# An organotypic *in vitro* model of matured blood vessels

**DOI:** 10.1101/2020.08.03.234807

**Authors:** Jaehyun Lee, Esak Lee

## Abstract

Angiogenesis is a physiological process in which brand-new blood vessels are formed from pre-existing blood vessels. The angiogenic processes are achieved by multiple steps, including angiogenic vascular sprouting, lumen formation, mural cell (e.g., smooth muscle cells) recruitment, and vessel stabilization by the mural cell coverage of the neovessels. Especially, mural cell recruitment to and coverage of the newly formed endothelium is a fundamental process to provide fully matured, functional blood vessels. Although investigation of the mural cell interactions with endothelial cells is crucial not only for better understanding of vascular physiology, but also for treating numerous vascular diseases, there has been a lack of three-dimensional (3D) *in vitro* models that recapitulate spontaneous processes of the vascular maturation. In this study, we describe an organotypic *in vitro* model that represents multi-step, spontaneous vascular maturation processes, which includes angiogenic vessel sprouting, smooth muscle cell (SMC) recruitment, and the SMC coverage of the neovessels. Using the system, we could spatiotemporally control vessel sprouting and vessel stabilization/maturation; and revealed an optimal condition that could reconstitute SMC-covered, matured blood vessels in 3D *in vitro*. We may provide a new platform for future mechanism studies of vascular interactions to mural cells and vessel maturation; and for pre-clinical screening and validation of therapeutic agent candidates for treating vascular diseases.

## 1. Introduction

Angiogenesis is defined as new blood vessel formation from pre-existing blood vasculatures [1]. Angiogenesis plays central roles in embryonic development and postnatal maintenance of blood vessels under certain conditions, such as wound healing, inflammation, hypoxia, and tissue regeneration. Angiogenesis is tightly regulated by distinct signals at different time points to provide structurally and functionally matured blood vessels. In the angiogenesis and vascular maturation processes, endothelial cells (ECs) and mural cells, such as smooth muscle cells (SMCs) undergo a series of multiple steps. Briefly, ECs in a pre-existing blood vessel depolarize and make new vascular sprouts in response to pro-angiogenic growth factor signals, such as vascular endothelial growth factor A (VEGF-A) and vascular endothelial growth factor receptor 2 (VEGFR-2) signaling [2]. The newly formed vessel sprouts proliferate, migrate, and further polarize to form a tubular lumen structure, enabling blood perfusion through the neovessels [3]. Next, interstitial mural cells are recruited to the lumenized neovessels in response to pro-maturation factors, such as platelet-derived growth factor BB (PDGF-BB) and platelet-derived growth factor receptor β (PDGFR-β) [4]; and Angiopoietin-1 (Ang-1) and Tie-2 signaling [5] to cover and stabilize the vessels [6].

Dysregulation of angiogenesis and vascular maturation broadly impacts human pathophysiology, which is involved in chronic inflammation, ischemic diseases, vascular leakage, thrombotic events, hyper-/hypotension, stoke, and cancer. Many researchers have tried to understand the spatiotemporal regulations of angiogenesis and vessel maturations to identify therapeutic targets to normalize blood vessels in the pathologic conditions. However, despite the numerous efforts, it is still not completely understood how the multicellular behaviors of ECs and mural cells are regulated under various conditions in the plethora of pro-angiogenic and pro-maturation factors.

Recent advances in microscopic imaging technologies have enabled us to observe fine structure of blood vasculatures *in vivo* [7–9], yet, it is still difficult to fully analyze the physical and morphological features of angiogenesis, due to limitations in controlling instrumental variables (e.g, soluble diffusible factors in surrounding microenvironment) and *in vivo* model variables (e.g., skin thickness, breathing motions, sexual differences); and difficulties in long-term live imaging under anesthesia [7–9]. By contrast, conventional angiogenesis assays *in vitro* in two-dimensional (2D) space in which ECs are cultured on plastic dishes or transwells are highly controllable [10], but are in a lack of relevant anatomies of three-dimensional (3D) vessel organization *in vivo*. For example, 2D cell culture models are often difficult to demonstrate directional sprouting of ECs in 3D extracellular matrix (ECM), vascular lumen formation, spatiotemporal mural cell recruitment/coverage, and the luminal flow along the EC-lining lumen [11]. Some 3D *in vitro* approaches, including bead sprouting assays and aortic ring explants [11] improved structural features in vessel sprouting and lumen formation, but still did demonstrate luminal flow and mural cell coverage on the vessels.

To address the issue, we develop an organotypic *in vitro* culture model that reconstitutes angiogenic sprouting and vessel maturation in 3D. We employ a polydimethylsiloxane (PDMS)-based microfluidic platform with two parallel microchannels fully embedded in collagen, where human umbilical vein endothelial cells (HUVECs) and vascular smooth muscle cells (SMCs) are separately seeded and cultured. Angiogenic and maturation factors are spatiotemporally introduced to facilitate HUVEC-SMC interactions in the middle of the two channels. The model allows us to screen maturation factors, such as PDGF-BB and Ang-1 and identify the optimal condition that creates matured blood vessels *in vitro*.

## 2. Materials and Methods

### 2.1. Cell culture

Human umbilical vein endothelial cells (HUVECs) were purchased from Lonza and cultured in EGM-2 (Lonza). Human vascular smooth muscle cells (SMCs) were purchased from Lonza were cultured in SmGM-2 (Lonza). All cells in the study were routinely tested to confirm no mycoplasma contamination. HUVECs were used at passages 3-8, and SMCs were used at passages 3-6. All the cells were maintained in standard tissue culture incubators at 37°C, 95% humidity, and 5% CO_2_.

### 2.2. Device fabrication

Our organotypic matured blood vessel on-chip was composed of two polydimethylsiloxane (PDMS) gaskets, cast from silicon wafer masters. The two gaskets were bonded after plasma etching, treated with 0.1 mg/ml poly-L-lysine (Sigma) for 3 hours and subsequently treated with 1% glutaraldehyde (EMS) for 15 min. Rat tail collagen I (Corning, 2.5 mg/ml) was pipetted into the devices after 2 acupuncture needles of 400 μm diameter were inserted in the device.

### 2.3. Co-culture of ECs/SMCs in the device

Once the collagen I was solidified, the casting needles were removed to leave two empty cylindrical channels completely embedded in the collagen I. Afterwards, HUVECs or SMCs were seeded into one channel through the reservoir (cell seeding density: HUVECs at 1 million cells/ml; SMCs at 0.3 million cells/ml) and allowed to form a monolayer cylindrical channel. Chemo-attractants have been introduced to the other channel to induce cell migration. For HUVEC sprouting we used MVPS cocktail (MVPS: MCP-1(75 ng/mL), VEGF-A (75 ng/mL), PMA (75 ng/mL), and S1P (500 nM) [12]) as previously described. For SMC migration, we used PDGF-BB (R & D systems) at 100 and 500 ng/ml. In some experiments, HUVECs and SMCs were seeded into two separate channels in the device at different time points to optimize cell migration of these cells toward the opposite direction and facilitate cellcell interactions between two cell types without hypertrophy. Cell-seeded microfluidic chip devices were placed on a platform rocker to generate gravity-driven luminal flow through the HUVEC and SMC channel at 37°C, 5% CO_2_, 95% humidity. The culture media and the chemotactic cytokines/growth factors in the device were replenished daily.

### 2.4. Immunofluorescence

Devices were fixed with 4% paraformaldehyde, permeated with PBST (1% Triton-X in PBS) at room temperature; then blocked with 3% bovine serum albumin (BSA) overnight at 4°C. Primary antibodies detecting alpha smooth muscle actin (αSMA, Abcam, rabbit, 1:100) and CD31 (Dako, mouse, 1:200) diluted in the blocking buffer (3% BSA) were treated into the devices and the devices were incubated overnight at 4°C. Primary antibodies were washed overnight using PBS at 4°C. Secondary antibodies (all from Invitrogen, 1:500) and DAPI (Millipore Sigma, 1:500) in the blocking buffer were subsequently treated into the devise and the devices were incubated overnight at 4°C. The devices were washed in PBS to remove fluorescent background before the confocal microscopy. Confocal images were acquired with a Leica SP8 Confocal microscope (Leica), then the images were adapted using the ImageJ (NIH).

### 2.5. Quantification and statistics

The statistical significance was tested using Mann–Whitney U-test. Error bars represent standard errors (SEM). All P values will be two-sided, and P < 0.05 will be considered statistically significant.

## 3. Results

### 3.1. Microfluidic device for reconstitution of 3D matured blood vasculatures in vitro

To reconstitute matured blood vessels 3D *in vitro*, we engineered an organotypic model of EC-SMC interactions based on a previously developed vessel-on-chip [12]. Briefly, our organotypic model is composed of two hollow cylindrical channels, which are fully embedded into 3D collagen I matrix (Figure 1a). In one of the channels, we seeded ECs (HUVECs) to form a biomimetic blood vessel. In a parallel channel, we seeded SMCs and allowed them to adhere to form a monolayer. We pursued for each type of cells sprouted toward the opposite channel in response to gradient of different growth factors. We hypothesized that if ECs and SMCs meet each other in the middle of the two channels; and the maturation factors were appropriately introduced, matured blood vessels are spontaneously formed in the middle of the two channels (Figure 1b).

**Figure 1.**
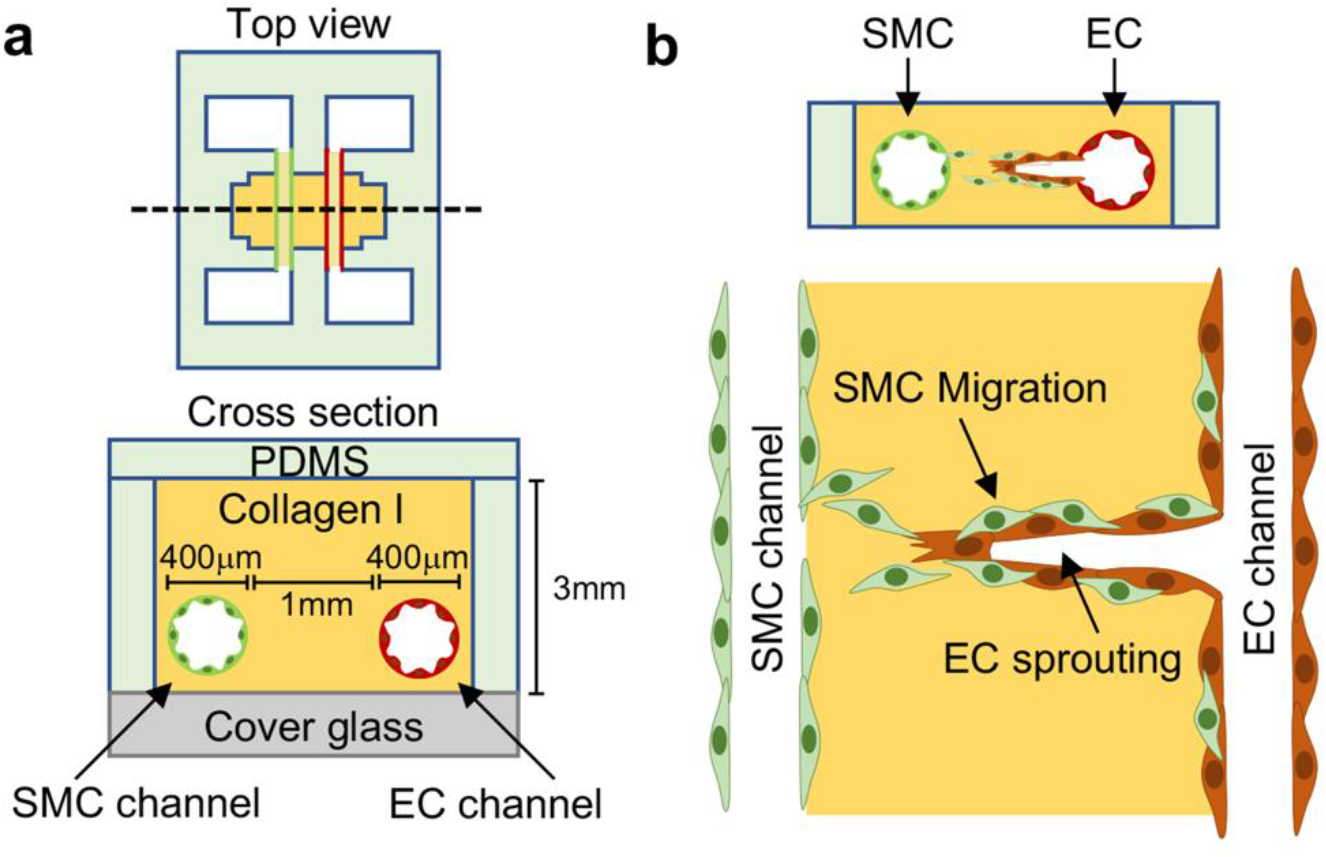
Schematic illustration of a matured blood vessel on-chip and expected cell dynamics of ECs and SMCs. (a) Device schematic. Polydimethylsiloxane (PDMS)-based microfluidic device with two parallel channels fully embedded in collagen, where SMCs and ECs are seeded separately. (b) Expected matured blood vessels in the middle of two channels with ECs and SMCs. ECs: endothelial cells, SMCs: smooth muscle cells.

### 3.2. SMC migration under different angiogenesis-related growth factors in vitro

Before seeding two cell types in one device, we first created a monoculature of SMCs, and examined the process of SMC migration in response to angiogenesis-related growth factors introduced in the opposite empty channel to form a chemotactic gradient (Figure 2). SMCs were seeded into one of the channels; and the MVPS cocktail (MVPS = monocyte chemotactic protein-1 (MCP-1), vascular endothelial growth factor-A (VEGF-A), phorbol 12-myristate 13-acetate (PMA), and sphingosine-1-phosphate (S1P) [12]) was added to the other channel (Figure 2a). We also evaluated platelet-derived growth factor-BB (PDGF-BB), a major stimulant for SMC migration and proliferation [4, 13] (Figure 2b-c). SMCs migration was monitored for next 3 days by taking bright-field images with differential interference contrast (DIC) microscope (Figure 2a-c). A previous study has screened the effects of various angiogenic factors and identified that MVPS cocktail induced the greatest EC sprouting with minimal single-cell migration of ECs in 3D *in vitro* [12]. In contrast to the EC behavior, SMCs did not invade considerably the collagen matrix in response to the gradient of MVPS cocktail (Figure 2a). Instead, SMCs invaded significantly the collagen matrix toward the opposite channel and extended their collective migration in response to the gradient of PDGF-BB (Figure 2b-c). To characterize the migration of SMCs, we defined the length and angle of SMC migration as shown in Figure 2d and analyzed them quantitatively. Interestingly, 100 ng/ml of PDGF-BB induced SMCs to have more and longer branches than 500 ng/ml of PDGF-BB (Figure 2e), although both 100 ng/ml and 500 ng/ml of PDGF-BB induced directional migration of SMCs toward the source channel (Figure 2f). Thus, we decided to perform the following experiments with 100 ng/ml of PDGF-BB for SMC migration and the MVPS only for EC sprouting in our co-culture system.

**Figure 2.**
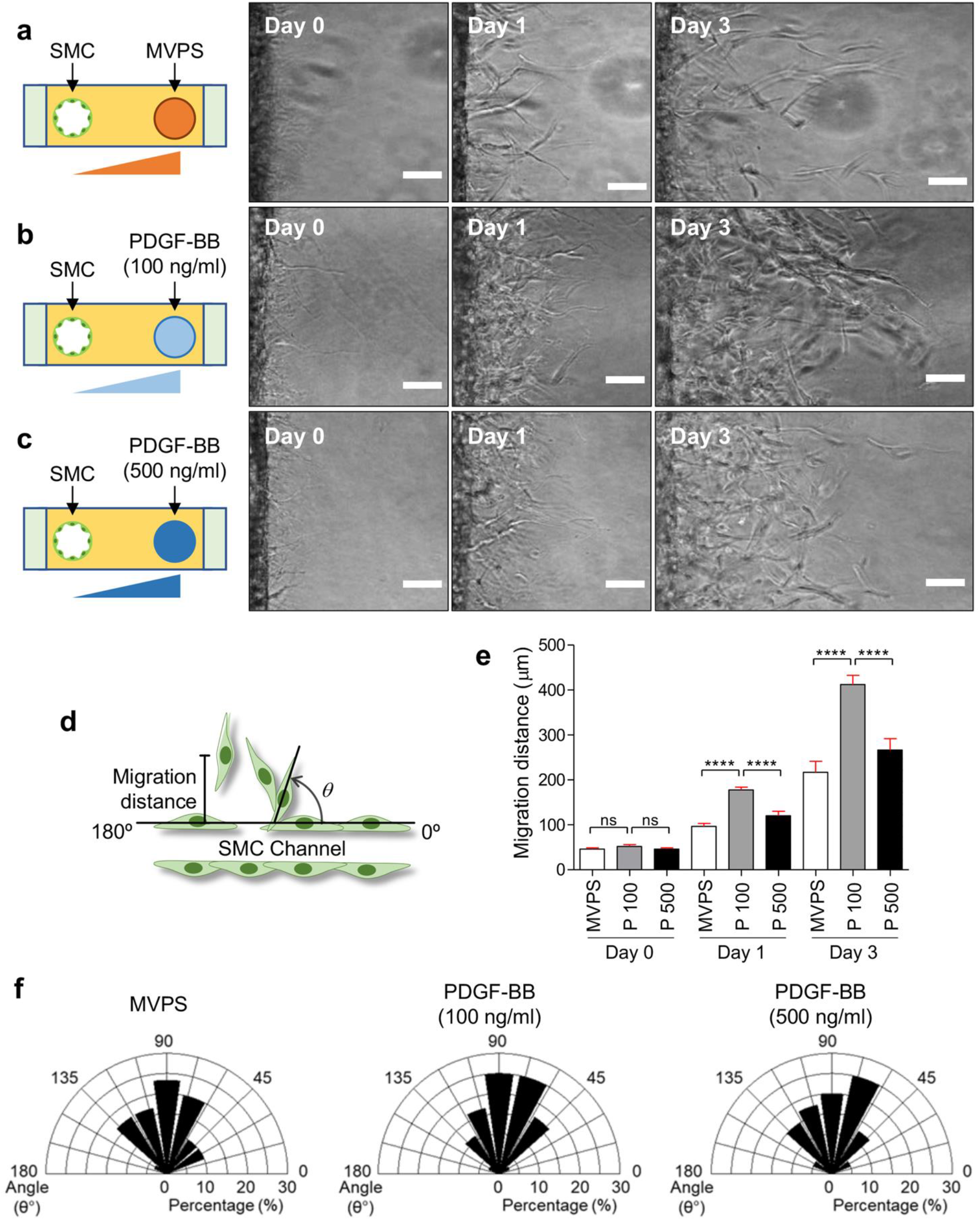
Characterization of SMC migration under angiogenesis-related growth factors. Representative time-lapse DIC images of SMCs exposed to MVPS (a), 100 ng/ml (b) and 500 ng/ml (c) of PDGF-BB at day 0, 1 and 3. Scale bars, 100 μm. (d) Definition of migration length and angle in SMC migration. Quantification of migration length (e) and angle (f) in SMC migration under MVPS, 100 ng/ml and 500 ng/ml of PDGF-BB. Mann–Whitney U-test **** p < 0.0001; ns, not significant.

### 3.3 The effect of co-stimulation of MVPS cocktail and PDGF-BB on SMC migration

Next, we sought to a way to stimulate ECs and SMCs in the co-culture system. Because MVPS cocktail was intended to promote EC sprouting [12] and the PDGF-BB was for the SMC migration, we examined the effect of co-stimulation of MVPS cocktail and PDGF-BB on SMC migration, before setting up the co-culture of ECs and SMCs (Figure 3). SMCs were seeded into one of the channels, and PDGF-BB 100 ng/ml was added to the opposite channel of SMCs as controls (Figure 3a). In the other group, SMCs were seeded into one of the channels; and PDGF-BB 100 ng/ml was added to the opposite channel of SMCs, while MVPS cocktail was added to the SMC channel at the same time (Figure 3b) to mimic future simulation of EC sprouting from the PDGF-BB channel in co-cultures (Figure 3b). SMC migration was monitored by taking cell images for next 4 days. For quantitative analysis, the angle between a migrating SMC and the SMC channel (θ); and the migration distance of the SMC from the channel were measured (Figure 3c-d). Compared to SMCs in the absence of MVPS cocktail, SMCs with MVPS showed limited migrations toward the PDGF-BB gradient (Figure 3c). In addition, SMC migration was more randomly oriented in presence of MVPS, while SMCs migration was nicely oriented to PDGF-BB source channel in the absence of MVPS cocktail (Figure 3d). These results indicate that SMC should not be stimulated simultaneously with both MVPS cocktail and PDGF-BB.

**Figure 3.**
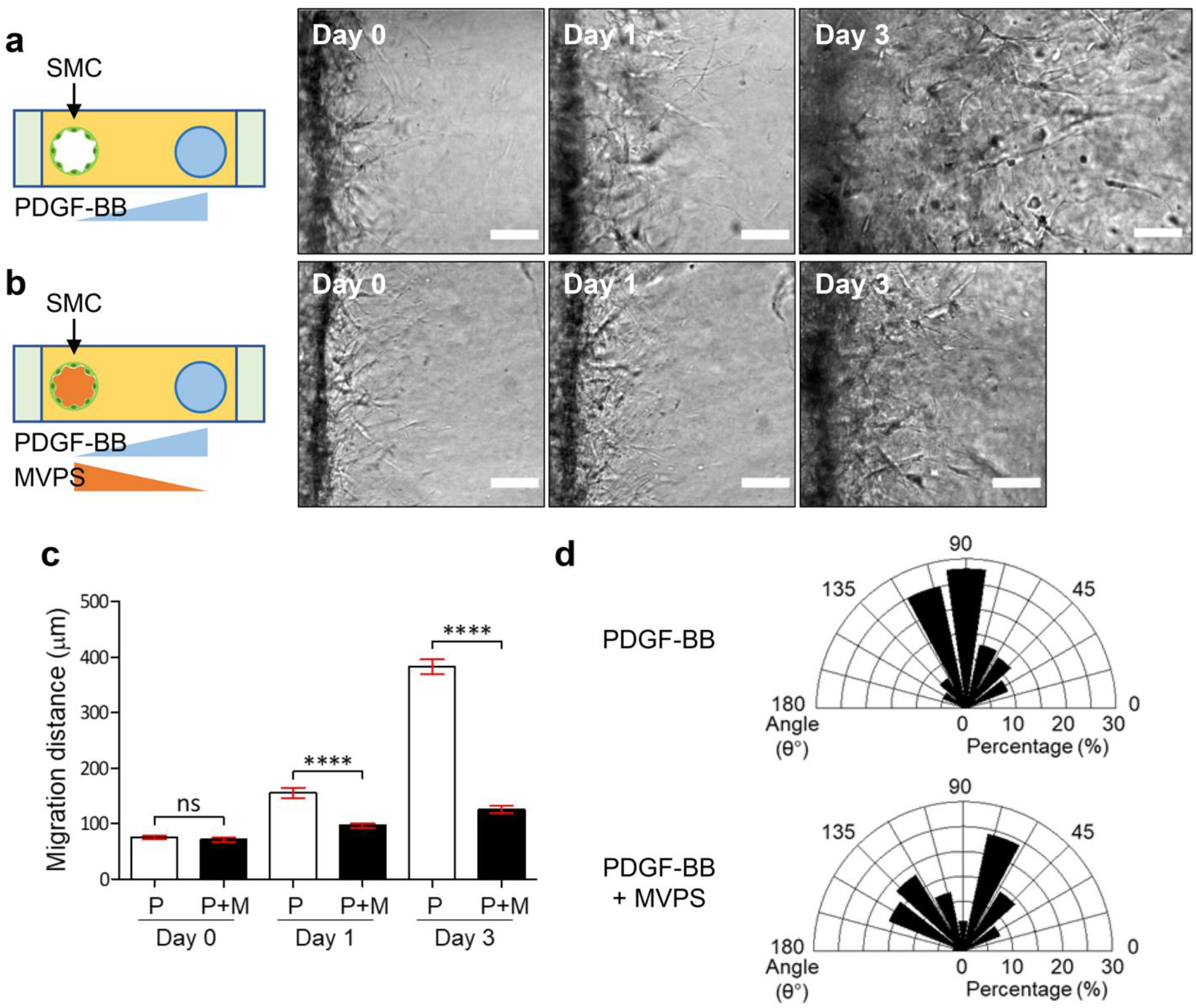
Characterization of SMC migration under co-stimulation of MVPS cocktail and PDGF-BB. Representative time-lapse DIC images of SMCs exposed to PDGF-BB alone (a) or both PDGF-BB and MVPS (b) at day 0, 1 and 3. Scale bars, 100 μm. Quantification of migration length (c) and angle (d) in SMC migration under PDGF-BB alone or both PDGF-BB and MVPS. Mann-Whitney U-test *** p < 0.001; ns, not significant.

### 3.4 Reconstitution of 3D matured blood vasculatures by spatiotemporal regulation of angiogenesis factors in co-culture of ECs and SMCs

To address the MVPS issue in SMC migration as well as the existing need for MVPS in EC sprouting in one system, we hypothesized that SMC migration with PDGF-BB needs to be preceded by EC sprouting with MVPS (Figure 4). Detailing the hypothesis, we expected to see robust EC sprouting in the monoculture of ECs with only MVPS (no SMC/PDGF-BB). After finishing the EC sprouting, MVPS would be removed. Then, SMCs would be seeded in the empty channel, then stimulated by PDGF-BB introduced in the EC channel, which may allow the robust induction of SMCs migration to the pre-made EC sprouts with only PDGF-BB. In the SMC induction step, we would use EGM-2 as a basal media to support EC survival without MVPS. To test the hypothesis, we co-cultured ECs and SMCs with a time difference of cell seeding and examined if EC sprouting or SMC migration fully supported the EC-SMC interactions and exhibited morphologic features of matured blood vessels (Figure 4). At day 0, only ECs were seeded in one of channels (right) and the MVPS cocktail was added to the other channel (left), then ECs were cultured for 3 days, allowing EC invasion and sprouting into collagen matrix (Figure 4a(i) and (ii)). At day 4, we rinsed the device with fresh EGM-2 to remove all the remaining MVPS cocktail before seeding SMCs. SMCs were additionally seeded into the left channel and 100 ng/ml PDGF-BB was added to the EC channel (right side) (Figure 4a(iii)). This is to mimic the nature of EC expression of PDGF-BB and the consequent recruitment of PDGFR-β expressing SMCs to the ECs [4, 13].

**Figure 4.**
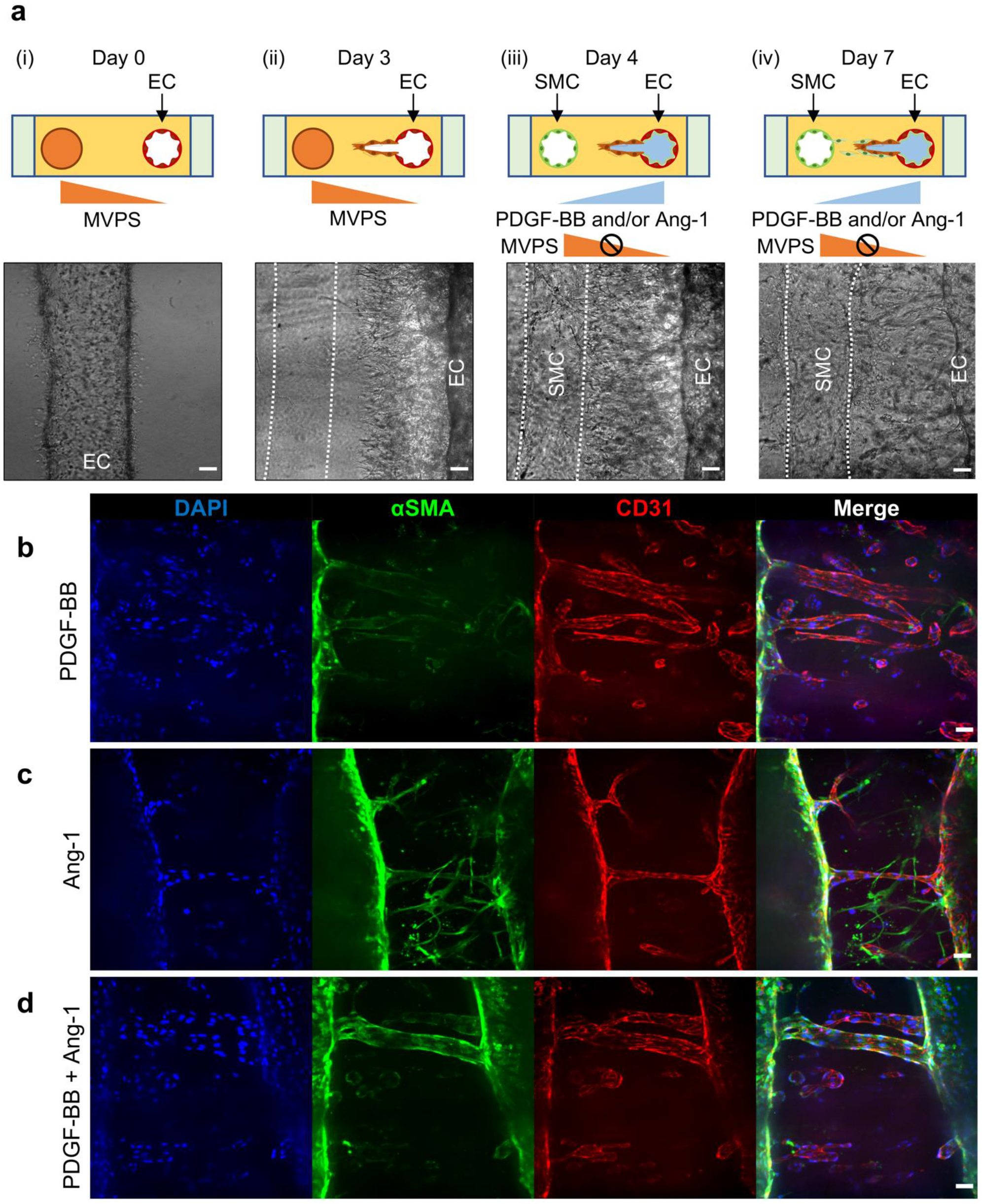
Reconstitution of 3D matured blood vessels by spatiotemporal regulation of angiogenesis-related factors. (a) Experimental set-up and representative time-lapse DIC image of microfluidic device at day 0 (i), 3(ii), 4(iii) and 7(iv). EC sprouting was induced by stimulation with MVPS for 3 days (i, ii). Then, SMC migration was induced by stimulation with PDGF-BB and/or Ang-1 for next 4 days, allowing the separate induction of ECs and SMCs in one system. Scale bars, 100 μm. Representative confocal immunofluorescence images of 3D matured blood vessels at day 7 in response to different combination of pro-maturation factors, PDGF-BB (b), Ang-1 (c) or both PDGF-BB and Ang-1 (d). Scale bars, 100 μm.

At day 7, samples were fixed and immunostained with anti-alpha smooth muscle actin (αSMA) antibodies and anti-CD31 antibodies to visualize SMCs and ECs, respectively (Figure 4a(iv) and 4b). We observed that stimulation of PDGF-BB was not enough to recruit SMC and cover ECs because of the slow migration of SMCs, demonstrated by few green cells in the bulk between the channels (Figure 4b). To overcome this, we additionally introduced 100 ng/ml of Angiopoietin-1 (Ang-1) to the EC channel, because it is reported that Ang-1 is critical for vascular homeostasis via promoting vascular maturation and integrity [5, 14]. We added the Ang-1 into the EC channel with or without PDGF-BB (Figure 4c-d) to show any synergistic effects. Stimulation with Ang-1 alone showed much faster migration of SMCs to the bulk compared than the single stimulation with PDGF-BB, reaching out to ECs channel but showed poor coverage (Figure 4c). Interestingly, co-stimulation of PDGF-BB and Ang-1 showed the highest SMC migration and coverage around the ECs (Figure 4d).

## 4. Discussion

Angiogenesis requires well-coordinated physiological stimulations with multiple growth factors to ECs and mural cells to generate matured, functional blood vessels [15, 16]. In the present study, we created an organotypic *in vitro* model to reconstitute matured blood vessels in 3D. We spatiotemporally controlled cell seeding (SMCs and ECs) and growth factor induction (MVPS, PDGF-BB, Ang-1) to provide neovessel sprouting and mural cell recruitment and coverages in 3D *in vitro*. There have been several *in vivo* and *in vitro* models to study the angiogenic processes [17–19]. Some *in vivo* models, including sponge/Matrigel assays [20], mouse retinopathy assays [21], fluorescent zebrafish assays [22, 23], and mouse tumor models [24, 25] have been developed and contributed to the vascular biology field. However, they have had limitations in spatiotemporal imaging and manipulation of the biological factors in the complex animal models, which hurdles more detailed, mechanistic understanding. In contrast, traditional *in vitro* models in 2D, such as scratch/wound healing assays [26], Boyden chamber/trans-well assays [18], stencil assays [27], and tube formation assays [28] are highly controllable, but poorly represent 3D multicellular organization of angiogenesis. Some 3D models including bead assays with spheroids composed of ECs and mural cells have been used [29, 30], showing 3D sprouting phenotypes, yet the model has limited imaging capacity inside the spheroids and the coverage was not fully achieved. Brudno et al. employed ECs and pericytes in the bead assays and showed EC sprouting and pericyte migration with nice imaging modalities [31], yet this model barely recapitulated perfusable 3D blood vessels with functional lumens.

Considering these limitations, in this study, we developed a 3D organotypic model using a PDMS-based microfluidic platform that recapitulated angiogenic sprouting, mural cell recruitment and mural cell coverage. Based on Nguyen et al. describing a 3D organotypic model that recapitulated morphogenesis of the angiogenic sprouting [12], we created two parallel channels fully embedded in 3D collagen matrix and cocultured ECs and SMCs separately in each channel. By introducing multiple growth factors in different time points to different directions, we have reconstituted vessel sprouting and vessel maturation in 3D. Compared to other models described above, our model was not only reproducible, capable of long-term experiments and imaging, but also recapitulating morphological features of luminal blood vasculature and their interactions to mural cells without exposing cells to artificial walls, such as silicone or glass, which affect cell behavior. Noteworthily, our microfluidic platform with two parallel channels allows us to spatiotemporally regulate and create gradients of different stimulating factors that affect EC-SMC interactions. Our results imply the importance of the spatiotemporal manipulation of growth factors formed different gradients during the vascular sprouting and maturation.

Angiogenesis consists of tightly regulated remodeling steps: i) EC sprouting from pre-existing blood vasculature, ii) EC tubule formation, acquiring a lumen, iii) mural cell recruitment, and iv) EC stabilization/vasculature maturation by mural cells [15, 16]. Each step requires proper angiogenesis-related factors. EC sprouting and tubule formation are governed by pro-angiogenesis factors, while EC stabilization/vasculature maturation is governed by pro-maturation factors. VEGF-A, the major pro-angiogenic factor, is known to be responsible for EC sprouting [32]. Like VEGF-A, MCP-1 promotes chemotaxis of ECs in a gradient of nanomolar concentrations [33]. In addition, Nguyen et al. reported that VEGF-A and MCP-1 alone did not promote substantial EC sprouting into the collagen matrix, while S1P and PMA alone promoted oriented EC sprouting into the collagen matrix toward the source channel [12]. They showed that a combination of MCP-1, VEGF, PMA, and S1P (the MVPS cocktail) resulted the greater EC sprouting response than single stimulation. Given the study, we induced EC sprouting towards the MVPS cocktail perfused channel (Figure 4a (i)–(ii)) [15, 16].

In the vessel maturation process, PDGF-BB is secreted by ECs and the ECs recruit PDGFR-β expressing SMCs, thus PDGF-BB is necessary for SMC migration and proliferation as a pro-maturation factor [4, 34, 35]. PDGF-BB is highly secreted by ECs, especially by tip cells of the newly formed sprouts to recruit mural cells eventually for EC stabilization/vasculature maturation [15, 16]. The absence of PDGF-BB causes deficiency of mural cell recruitment around ECs, and results in abnormally dilated capillaries and reduced deposition of basement membranes. Inspired by these phenomena in nature, we firstly induced EC sprouting using the MVPS cocktail, then seeded SMCs in the empty channel where MVPS was previously. Next, we introduced PDGF-BB at different concentration (100 ng/ml and 500 ng/ml) in the EC channel, to form the gradient of PDGF-BB across the collagen matrix. Interestingly, 100 ng/ml of PDGF-BB stimulation showed robust migration of SMCs than 500 ng/ml of PDGF-BB stimulation toward the PDGF-BB source channel. Previous study reported that PDGF-BB establishes a negative feedback loop by controlling PDGF-BB receptor expression via c-Myc [36, 37]. In our experiments, it is likely that 500 ng/ml of PDGF-BB was an overtreatment, saturating the entire system with excess PDGF-BB, which caused less migration of SMCs than 100 ng/ml of PDGF-BB treatment (Figure 2b-c). As comparison to PDGF-BB, the gradient of MVPS cocktail did not induce significant invasion of SMCs into the collagen matrix (Figure 2a).

Interestingly, SMCs did not invade actively the collagen matrix when they were exposed to both the gradient of PDGF-BB from the opposite channel and direct perfusion of MVPS cocktail via SMC channel, simultaneously (Figure 3). This result might be related to a previous study showing that VEGF-A mediated activation of VEGFR-2 suppressed PDGFR-β signaling in SMCs by forming a non-canonical PDGFR-β/VEGFR-2 receptor complex, which limited the PDGFR-β signal transduction [38]. Taken together, our results emphasize the importance of spatiotemporal control of growth factors in blood vascular engineering.

In co-culture of ECs and SMCs with a delay of seeding SMCs, we observed that under single stimulation of PDGF-BB, SMCs did not cover EC sprouts sufficiently and few SMCs migrated into the collagen matrix between the channels (Figure 4b). Under single stimulation of Ang-1, SMCs migrated well and reached the parent EC channel, but poorly covered EC sprouts (Figure 4c). In addition to PDGF-BB, Ang-1 is also a crucial pro-maturation factor. It is known that Ang-1 decreases basal permeability of ECs by strengthening adhesion and junction molecules including platelet endothelial cell adhesion molecule-1(PECAM-1) and vascular endothelial cadherin (VE-cadherin) junctions, and mediates EC survival and mural cell attachment by attenuating VEGF-A mediated angiogenic sprouting [39–42]. Apart from PDGF-BB secreted by ECs, Ang-1 is secreted by mural cells and recognized by Tie-2 receptor expressed by ECs [15, 16]. Previous study has reported that the cell-cell adhesion molecule, such as N-cadherin between EC and mural cells, are required for rapid migration of SMCs [43]. Such cellular contact migration guidance is commonly used in leukocyte trafficking to endothelium for fast arrival to the inflamed sites [44–46]. Similarly, it is likely that EC-SMC interaction via N-cadherin and/or other adhesion molecules that are mediated by Ang-1, would give an advantage in the fast migration of SMCs. Another previous study revealed that SMCs can migrate directionally without an exogenously established chemotactic gradient [47]. Co-stimulation of PDGF-BB and Ang-1 showed that highest coverage of SMCs around the ECs sprouts as well as parent EC channel, suggesting synergic effect of PDGF-BB and Ang-1 (Figure 4d). It is also likely that co-stimulation of PDGF-BB and Ang-1 may encourage collective migration of SMCs, presenting robust coverage of ECs. Taken together, we identified the role of PDGF-BB and Ang-1 in SMC migration and EC coverage and established the optimal condition that generated matured blood vessels with SMC coverage *in vitro*. Further validations with these growth factor combinations for matured blood vessels *in vivo* remain for future studies.

Dysfunctional EC-SMC interactions have been involved in a variety of congenital and acquired diseases [48]. For example, idiopathic basal ganglia calcification is caused by a loss of function mutation in PDGF-BB [49]. Molecules in EC-SMC interactions have been recognized as therapeutic targets, inhibiting angiogenesis in tumor microenvironments [50, 51]. For example, Trebananib, an Ang-1/-2 neutralizing petibody, inhibits Ang-1/Tie-2 interactions [52, 53]. Receptor tyrosine kinase inhibitors, such as Cediranib, Nintedanib and Sorafenib inhibit signaling pathways of VEGFRs and PDGFRs [54]. Since our *in vitro* model well recapitulates the process of angiogenesis from EC sprouting to maturation by SMCs, our model may provide a unique platform for drug screening and mechanism studies. Further studies with modifications of the current model may focus on more specific blood vascular diseases or lymphatic endothelial cell (LEC) interaction with SMCs to identify molecular mechanisms and cellular behaviors involved in blood and lymphatic vessel related diseases.

## 5. Conclusion

In conclusion, we describe a microfluidic platform that reconstitutes the complex process of angiogenesis and vessel maturation in 3D *in vitro*. We reveal that spatiotemporal regulation of stimulus factors is critical to generate EC sprouts and SMC migration simultaneously in one system and that co-stimulation with PDGF-BB and Ang-1 creates matured blood vessels than a singular stimulation. The key features of our model are the followings: i) a microfluidic platform with two separate channels, which allows us not only to create the gradient of variable factors but also to control variable factors spatiotemporally, ii) the ability to be prepared with laboratory equipment and easy handling, iii) the capability of quantitatively analyzing cell dynamics, iv) the potential applicability to lymph-angiogenesis or other blood vascular disease models. We suggest that our microfluidic chip could be used for screening and pre-clinical validation of drug candidates for vascular diseases as well as the investigation of fundamental angiogenesis studies.

## Acknowledgements

J.L. and E.L. are supported by the Cornell University Startup funds, and the Nancy and Peter Meinig Family Investigator funds. This work was performed in part at the Cornell NanoScale Facility (CNF), a member of the National Nanotechnology Coordinated Infrastructure (NNCI), which is supported by the National Science Foundation (Grant NNCI-1542081).

